# Evaluating DNA methylation age on the Illumina’s methylationEPIC BeadChip

**DOI:** 10.1101/466045

**Authors:** Dhingra Radhika, Lydia Coulter Kwee, David Diaz-Sanchez, Robert B. Devlin, Wayne Cascio, Carol Haynes, Elizabeth R. Hauser, Simon Gregory, Svati Shah, William Kraus, Kenneth Olden, Cavin K. Ward-Caviness

**Affiliations:** National Health and Environmental Effects Laboratory, US Environmental Protection Agency, Chapel Hill, NC, USA; Department of Environmental Sciences and Engineering, Gillings School of Public Health, University of North Carolina, Chapel Hill, NC, USA; Institute for Environmental Health Solutions, University of North Carolina, Chapel Hill, NC USA; Duke Molecular Physiology Institute, Duke University Medical Center, Durham, NC, USA; Department of Biostatistics and Bioinformatics, Duke University Medical Center, Durham, NC, USA; Cooperative Studies Program Epidemiology Center, Durham Veterans Affairs Medical Center, Durham, NC, USA; Division of Cardiology, Department of Medicine, School of Medicine, Duke University, Durham, NC, USA; National Center for Environmental Assessment, US Environmental Protection Agency, Chapel Hill, NC, USA

## Abstract

DNA methylation age (DNAm age) has become a widely utilized epigenetic biomarker for the aging process. The Horvath method for determining DNAm age is perhaps the most widely utilized and validated DNA methylation age assessment measure. Horvath DNAm age is calculated based on methylation measurements at 353 loci which were present on Illumina’s 450k and 27k DNA methylation microarrays. With the increasing use of the more recently developed Illumina MethylationEPIC (850k) microarray, it is worth revisiting this widely used aging measure to evaluate differences in DNA methylation age estimation based on array design. Of the requisite 353 loci, 17 are missing from the current 850k microarray. Using 17 datasets with 27k, 450k, and/or 850k methylation data, we calculated and compared each sample’s epigenetic age estimated from all 353 loci required from the Horvath DNAm age calculator (full), and using only the 336 loci present on the 27k, 450k, and 850k arrays (reduced). In 450k/27k data, missing loci caused underestimation of epigenetic age when compared with the full clock. Underestimation of full epigenetic age grew from ages 0 to ~20, remaining stable thereafter (mean= −3.46 y, SD=1.13) years for individuals ≥20 years. Underestimation of DNAm age by the reduced 450k/27k data was similar to the underestimation observed in the 850k data indicating that array differences in DNAm age estimation are primarily driven by missing probes. Correlations between age and DNAm age were not dependent on missing probes or on array designs and consequently associations between DNAm age and outcomes such as sex remained the same independent of missing probes and probe design. In conclusion, DNAm age estimations are array dependent driven by missing probes between arrays. Though correlations and associations with DNAm age may remain the same, researchers should exercise caution when interpreting results based on absolute differences in DNAm age or when mixing samples assayed on different arrays.

**Disclaimer:** This paper has been reviewed by the National Health and Environmental Effects Research Laboratory, U.S. EPA, and is approved for publication. Approval does not signify that the contents necessarily reflect the views and policies of the agency, nor does mention of trade names or commercial products constitute endorsement or recommendation for use. The authors declare they have no competing financial interests.

## Introduction

DNA methylation has recently shown promise as a potentially clinically useful biomarker of aging. A recent “epigenetic clock” developed by Horvath (1) has been shown to be an accurate estimator of age across multiple tissues and populations, and differences between DNA methylation age and chronological age are associated with pathophysiological biomarkers and incident disease (2).

The method developed by developed by Horvath (1) is perhaps the most widely used and validated epigenetic age estimation method; it relies on measurement of percent methylation at 353 loci (CpGs) on either the Illumina 450k (450k) or Illumina 27k (27k) microarray chips. Recently, Illumina released the Infinium MethylationEPIC Bead Chip (850k), which uses the same technology as the Illumina 450K microarray to assay 866,836 CpGs (3). Though the 850k microarray assays more loci, 8.9% of CpGs included on 450K microarray were omitted from the 850k microarray. In particular, 17 of the 353 CpGs (4.8%) necessary to calculate epigenetic age via the Horvath method are missing. While missing CpGs are imputed in the online calculator (4) to allow for estimation of epigenetic age, these missing probes may systematically bias the estimation of DNA methylation age and consequently alter the detection or interpretation of associations with health outcomes and inhibit cross-platform comparisons and analyses.

To evaluate the impact of microarray design changes on the estimation of DNA methylation age, we compared the Horvath DNA methylation age (DNAm age) calculated using all 353 CpGs (full DNAm age) to estimates obtained from using either the 27k or 450k platform while restricting to the 336 CpGs available on the 850k platform. We used 15 publicly available non-cancer blood tissue datasets (available in the Gene Expression Omnibus(GEO), https://www.ncbi.nlm.nih.gov/geo/), as well as blood samples from a cardiac catheterization cohort (CATHeterization GENetics; CATHGEN) where DNA methylation was assessed on both the 450k and 850k arrays.

## Methods

### Missing loci and datasets

To determine which loci in Horvath’s original epigenetic clock loci are missing from the 850k platform we compared the 850k manifest of probe loci and the list of loci required for Horvath’s estimation of epigenetic age (available in Additional File 3 of (1)).

From the 81 datasets used to develop the Horvath epigenetic clock, we selected those 15 datasets (detailed in Supp. Table 1) whose non-cancerous samples were drawn from blood (excluding cord blood), were publicly available on the Gene Expression Omnibus (GEO; https://www.ncbi.nlm.nih.gov/geo/) and whose methylation beta values were readily available on GEO. Though chronological age was not available in GSE42865 and GSE35069, and sex was not available in GSE30870 and GSE 42865, these datasets were also included in analyses that did not require age or sex.

Samples (N = 3,672) in the 15 eligible GEO datasets (summarized in Table S1) were drawn from people ages 0 to 101, and included whole blood, peripheral blood monocytes (PBMC) and single leukocyte cell types. GSE 19711 was divided into two datasets (controls and ovarian cancer cases) for consistency with the Horvath epigenetic clock manuscript (1). Though a few of these datasets include samples from cancer patients, the tissue obtained was non-cancerous, and their methylation age had previously shown no association to cancer (1). Further information about these datasets may be found on GEO, and in Additional file 2 of Horvath’s manuscript which describes these datasets and their rationale for inclusion in the development of his epigenetic clock (1).

In addition to the GEO datasets, two datasets from the Catheterization Genetics cohort (CATHGEN) were employed to compare the 450k and 850k platforms. CATHGEN participants were recruited from subjects undergoing an outpatient cardiac catheterization at Duke University from 2001-2011 (5). Ethics approval was administered by the Duke Institutional Review Board for CATHGEN.

The samples were processed by reading in the idat files using minfi v1.21.1, examining samples for exclusion based on Illumina’s default quality control (QC) procedures, background correction via minfi’s ssNoob, and extracting the un-normalized beta values. The CATHGEN samples processed on the 450k and 850k microarrays were not obtained from the same individuals, and no samples were excluded based on QC for the 450k microarray, while two samples from the 850k microarray were excluded. This left 205 CATHGEN samples for the 450k microarray (ages 23-91 y) and 568 samples available from the 850k microarray (ages 33-87 y).

### DNAm age processing

Methylation beta values were extracted from the downloaded GEO datasets, and were not further normalized before uploading to the (online) DNA Methylation Age Calculator as recommended (https://dnamage.genetics.ucla.edu/). Where GEO datasets were previously normalized, we deselected the normalize data option during processing in the DNA methylation calculator; otherwise, the normalize data option was selected for unnormalized data.

All samples were included from the publicly available data. Sex, age, sample id and blood type were extracted from the downloaded GEO datasets. The online DNA methylation age calculator automatically imputes any missing probes (https://labs.genetics.ucla.edu/horvath/dnamage/).

### The epigenetic clock across the age ranges in 450k/27k data

To ascertain how the 17 missing loci might systematically misestimate epigenetic age via Horvath’s 353-probe DNA methylation clock, we calculated DNA methylation age in 27k and 450k datasets (GEO & CATHGEN 450K datasets) with and without the 17 probes unavailable on the 850k microarray. For each GEO dataset, as well as the CATHGEN 450k datasets, DNAm age calculated using the reduced 450k data were compared to DNAm age calculated using the full 450k data, graphically and using summary statistics. The comparisons were repeated in subjects chronologically aged 20 y or less, and in ages ≥ 20 y, a cutoff selected based on the observed inflection point in the plot of age vs the difference in DNA methylation age estimated using the full and reduced 450k data.

We hypothesized that the relationship of DNA methylation age to chronological age differed in the full and reduced 450k/27k datasets and that the difference varied by chronological age group (>20 years and ≤ 20 years). Using all samples within each age group, we separately regressed full 450k DNAm age and the reduced 450k DNAm age on chronological age, and compared resulting the intercepts and chronological age slopes estimates. This analysis excluded the GSE42865 and GSE35069 datasets as chronological age was not publicly available.

Within each age group, we also assessed the possibility that the relationship between DNA methylation age, and thus age acceleration, and a clinical or other variable of interest could be modified by the loss of 17 missing loci from the dataset. As sex was the only widely available variable in the public data, we separately regressed age acceleration estimated based on the full and reduced 450k data on sex (ref. = Male), using all available samples within each age group. We repeated these analyses in each individual dataset, without regard to the chronological age of samples. We then statistically compared the slope obtained when using full 450k data age acceleration to that obtained via reduced 450k data age acceleration for models of the association of sex with age acceleration. Additionally, we compared residual plots of *full* and *reduced* 450k data DNAm age acceleration regressed on chronological age for all GEO datasets where age was available in the CATHGEN 450k dataset.

### Comparison of DNA methylation age in 450k and 850k datasets

The CATHGEN data were used to ascertain if technological changes in the 850k platform as compared to the 450k or 27k platforms contribute to mis-estimation of epigenetic age. To that end, *full* and *reduced* datasets for the samples processed on the 450k, as well as a dataset for the samples processed on the 850k were created for CATHGEN. Linear fits of the epigenetic age by chronological age for each of the 3 CATHGEN datasets were produced. The intercept and slopes of these linear fits were compared, to ascertain if the 850k platform impacts the methylation measurement such that it would impact the calculation of epigenetic age, in a manner separate from the effect of the 17 missing probes.

The CATHGEN dataset affords the ability to quantify any deviation of 850k DNAm ages from expected values. As no ‘correct’ estimate of DNAm age on the 850k is available, we chose regressed DNAm age on categorical variables for dataset types (full 450k and 850k in one model and reduced 450k and 850k in the second model) while controlling for age. In both models, the 450k DNAm age, either full or reduced” was the referent category.

### Software and statistical analyses

All work to determine the lost loci, to prepare data for the online DNA Methylation Age Calculator (https://dnamage.genetics.ucla.edu/) and to subsequently compare epigenetic age estimates with chronological age were performed in R (version 3.4.0) (6).

### Terminology

Three categories of DNA methylation data were used in this analysis: 1) data from the Illumina 450k array or the 27k array (“full 450k data”); 2) data from the Illumina 450k or 27k arrays with the 17 probes not on the Illumina 850k array removed (“reduced 450k data); and 3) data from the Illumina 850k array (“850k data”). “Reduced 450k DNAm age” and “full 450k DNAm age” refer to the application of the Horvath epigenetic clock to reduced and full 450k data, respectively.

## Results

### Missing probes & descriptions of the datasets

The 17 required DNA methylation age loci that are not included in the 850k manifest are listed in Table 1. The GEO and CATHGEN 450k datasets together encompass 3,973 individuals (52% female, among those reporting sex) whose ages range from 0 (i.e., newborn) to 101 years (Table 2). In addition, we had 568 independent CATHGEN samples that were processed on the 850k platform.

**Table 1.** Missing probes, SNP presence, and symbol.

| CpG | SNP? | Symbol |
| --- | --- | --- |
| cg19945840 | no | B3GALT6 |
| cg02972551 | no | JMJD1A |
| cg02654291 | yes | C9orf64 |
| cg13682722 | yes | C14orf102 |
| cg09869858 | yes | P11 |
| cg06117855 | yes | CLEC3B |
| cg05590257 | yes | LOC201164 |
| cg27016307 | yes | HRC |
| cg24471894 | yes | KIAA0020 |
| cg04431054 | no | LOC133619 |
| cg16494477 | no | FGF18 |
| cg19046959 | no | COL8A2 |
| cg17408647 | yes | FLJ10803 |
| cg27319898 | no | FLJ32110 |
| cg19569684 | no | PACAP |
| cg19273182 | no | PAPOLG |
| cg09785172 | no | WFS1 |

**Table 2.** Comparison of DNA methylation age (DNAm age) estimation from full 450k data, reduced 450k data, and 850k data in GEO and CATHGEN datasets. The mean, standard deviations and correlation with chronological age (Age corr.) of DNAm age are provided for each dataset.

| GEO Series no. | Platform | N (prop. female) | Chronological age |  | (Full) 450k data (353 loci) |  | Reduced 450k or 850k data (336 loci) |  | Comparison |
| --- | --- | --- | --- | --- | --- | --- | --- | --- | --- |
|  |  |  | Median (range) | Mean (SD) | Mean (SD) | Age corr. | Mean (SD) | Age corr. | (450k data DNAm age) - (red. 450k data DNAm age) |
| GSE19711cases (7,8) | 27K | 266 (1.0) | 67 (49, 91) | 66.42 (9.35) | 62.5 (11.47) | 0.55 | 58.43 (11.02) | 0.56 | 4.11 (0.81) |
| GSE19711controls (7,8) | 27K | 274 (1.0) | 64 (52, 78) | 64.89 (6.74) | 62.57 (7.65) | 0.66 | 58.56 (7.52) | 0.66 | 4.01 (0.68) |
| GSE20067 (7,9) | 27K | 192 (0.51) | 43 (24, 74) | 43.9 (9.8) | 43.45 (9.27) | 0.81 | 38.55 (9.2) | 0.81 | 4.85 (0.95) |
| GSE20236 (10) | 27K | 93 (1.0) | 63 (49, 74) | 62.86 (6.33) | 53.79 (6.51) | 0.69 | 49.92 (6.32) | 0.68 | 3.87 (0.58) |
| GSE20242 (10) | 27K | 50 (0.74) | 34 (16, 69) | 35.86 (13.89) | 45.02 (27.45) | 0.55 | 41.49 (27.71) | 0.53 | 2.30 (0.84) |
| GSE27097 (11) | 27K | 398 (0.0) | 9.3 (3.6, 17.8) | 9.89 (3.63) | 9.6 (4.41) | 0.75 | 8.14 (3.88) | 0.72 | 1.46 (0.69) |
| GSE30870 (12)** | 450K | 38 (0.74) | 44.5 (0, 100) | 46.32 (47.01) | 41.06 (42.02) | 0.99 | 38.93 (39.95) | 0.99 | 2.14 (2.13) |
| GSE32149 (13) | 450K | 48 (0.52) | 15 (3.5, 76) | 22.15 (18.43) | 22.3 (15.13) | 0.96 | 19.96 (14.34) | 0.97 | 2.34 (0.92) |
| GSE35069 (14)* | 450K | 60 (0.0) | NA | NA | 41.74 (12.75) | - | 39.15 (12.84) | - | 2.59 (0.56) |
| GSE36064 (11) | 450K | 78 (0.0) | 3.1 (1.0, 16.1) | 4.58 (4.11) | 4.38 (3.92) | 0.93 | 3.62 (3.27) | 0.93 | 0.76 (0.66) |
| GSE40279 (15) | 450K | 656 (0.52) | 65 (19, 101) | 64.04 (14.74) | 63.08 (11.53) | 0.91 | 60.67 (11.66) | 0.92 | 2.41 (0.70) |
| GSE41037 (16) | 27K | 720 (0.38) | 33 (16, 88) | 37.4 (15.72) | 36.85 (15.38) | 0.95 | 33.07 (15.07) | 0.96 | 3.81 (0.79) |
| GSE41169 (16) | 450K | 95 (0.29) | 29 (18, 65) | 31.57 (10.28) | 31.23 (11.01) | 0.94 | 27.67 (10.69) | 0.94 | 3.55 (0.60) |
| GSE42861 (17) | 450K | 689 (0.71) | 54 (18, 70) | 51.93 (11.8) | 53.38 (11.09) | 0.90 | 50.22 (11.01) | 0.90 | 3.16 (0.58) |
| GSE42865 (18)* ** | 450K | 15 (0.62) | NA | NA | 38.19 (9.45) | - | 35.68 (9.68) | - | 2.40 (1.10) |
| CATHGEN 450k ° | 450k | 206 (0.37) | 64 (33, 87) | 63.41 (11.85) | 64.58 (10.50) | 0.88 | 60.73 (10.23) | 0.87 | 3.85 (0.72) |
| CATHGEN 850k † ° | 850k | 568 (0.41) | 59 (23, 91) | 60.11 (12.44) | - | - | 58.16 (10.51) | 0.86 | - |

\* As chronological age was missing for these datasets, correlation with age and age acceleration could not be determined.
\*\* Proportion Female was obtained from supplemental table of the original epigenetic clock manuscript (Horvath, 2013), and were not available in GEO.
† Because the 17 loci required to complete the epigenetic clock are unavailable on the 850k platform, there is not information for the full epigenetic clock
° CATHGEN 450k and CATHGEN 850k are not comprised of the same individuals. That is, the underlying sample population is non-overlapping.

### Comparison of DNA methylation age in 450k data and 850k data

DNAm age estimated separately in the CATHGEN’s *full* 450k, *reduced* 450k and 850k datasets using the epigenetic clock all showed positive correlations with chronological age (Table 2, Figure 1). For each of these three datasets, the slope between DNAm age and chronological age is nearly identical (0.73-0.78). However, in a regression of DNAm age on dataset type (full 450k vs. 850k) correcting for age, 850k DNAm ages had a mean difference of - 3.96 y (95%CI: −4.08, −3.12; p <0.0001) as compared to the full 450k, which is very close to the underestimation seen with the when comparing CATHGEN DNAm age estimates from the reduced 450k data with the full 450k data (paired t-test: 3.85 y, p<0.0001). There was no significant difference between the 850k DNAm age and reduced 450k DNAm age in CATHGEN (−0.14; 95%CI: −0.98, 0.70, p=0.75).

**Figure 1.**
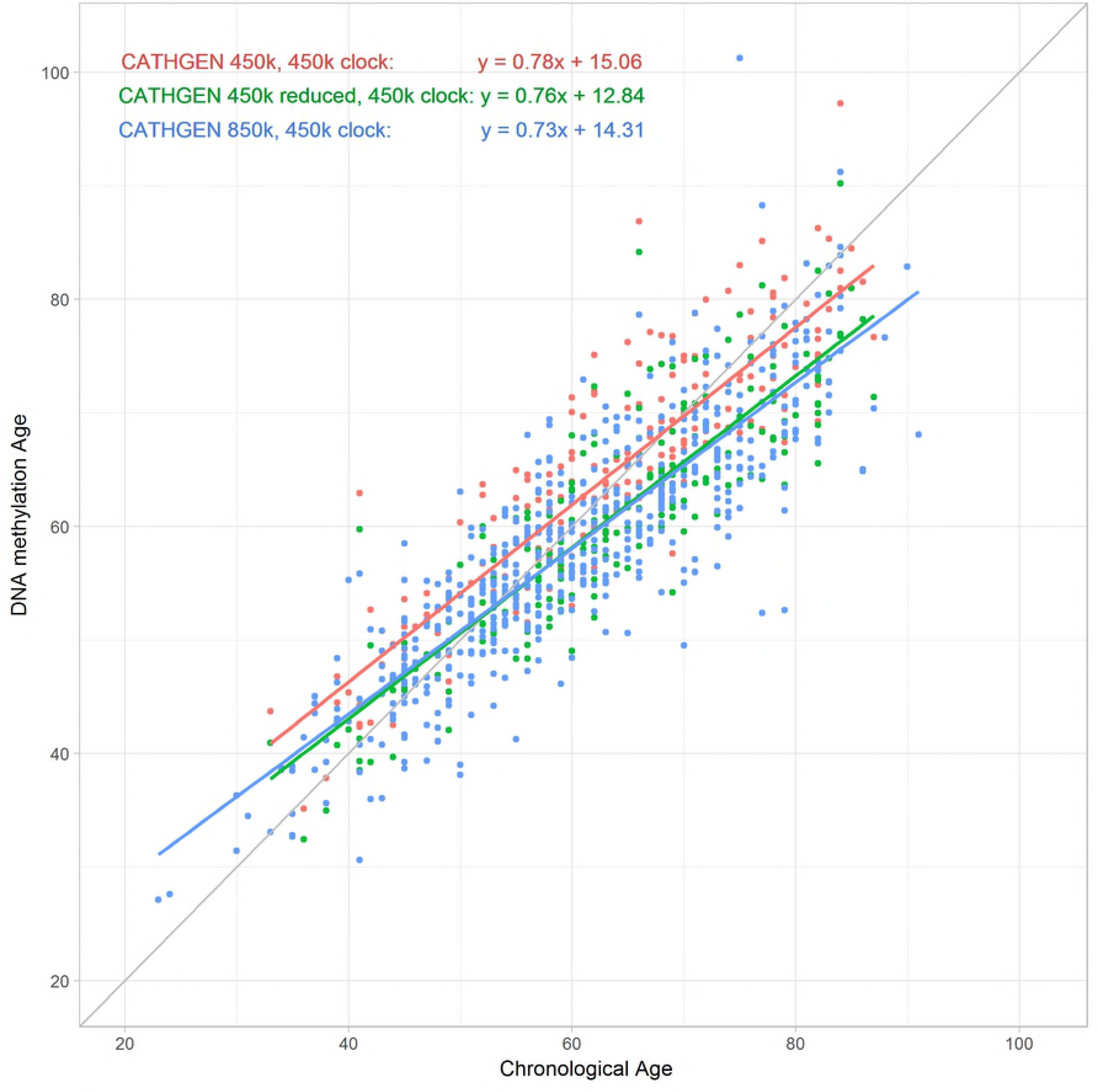
Epigenetic age by chronological age in combinations of CATHGEN dataset and epigenetic clock: The plot of DNA methylation by chronological age shows the impact of the 17 missing probes, by applying the epigenetic clock to CATHGEN 450k (‘full’ and ‘reduced’) and 850k datasets.

### Probe exclusion effects on Horvath DNAm age in 16 datasets

Across all 16 datasets with 450k or 27k data, reduced 450k DNAm age underestimated DNA methylation age as compared to the full 450k DNAm age (Figure S1). In peripheral blood samples from the youngest individuals (chronological age < 20 y), the individual difference between epigenetic age as estimated using the *full* and *reduced* datasets increased with age (Figure 2, Table 3). However, in samples from older individuals, (chronological age ≥ 20 y), the difference did not increase with age but we observed greater inter-individual variability in the difference between full and reduced DNAm age in older individuals (SD = 1.13) than in the younger age group (ages 0-5y: SD = 0.27; ages 5-10y: 0.35; ages 10-15y: SD= 0.54; and ages 15-20y: SD=0.82). Across all datasets, the correlation between full and reduced 450k data remained high ranging from 0.989 to 0.999.

**Figure 2.**
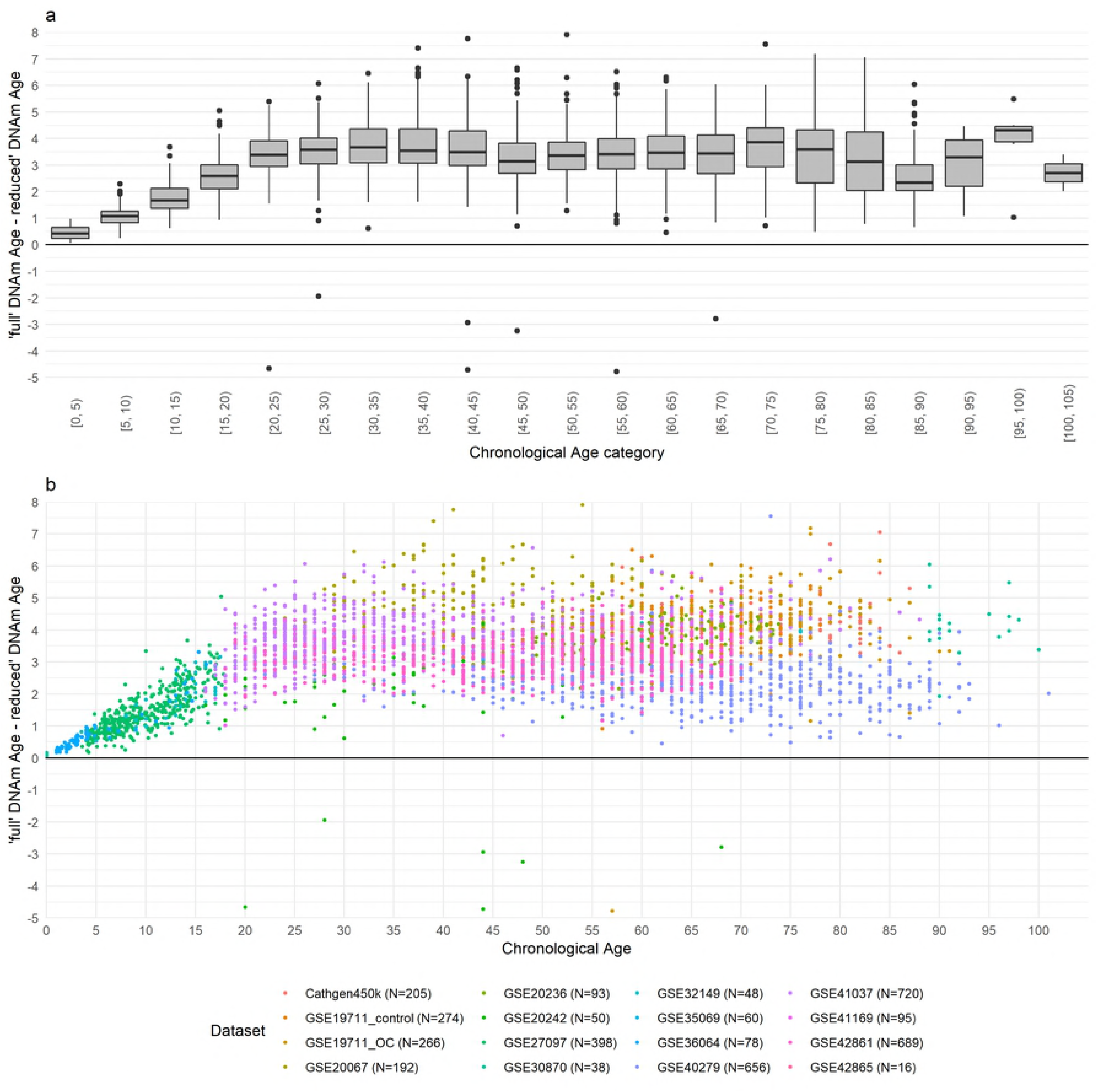
Difference of ‘full’ and ‘reduced’ epigenetic Age by chronological age. The difference of ‘full and reduced’ epigenetic ages calculated in the GEO (450k and 27k) and CATHGEN 450k data are presented as (a) boxplot by 5 year chronological age categories and (b) as a scatterplot.

**Table 3.** Regression of DNA methylation age on chronological age, by age group, in the full and reduced 450k/27k datasets (GEO and CATHGEN).

| Data | <u>Age < 20 years (N = 616)</u> |  | <u>Age ≥ 20 years (N =2,972)</u> |  |
| --- | --- | --- | --- | --- |
|  | Intercept | Chronological Age | Intercept | Chronological Age |
|  | Estimate (95% CI) | Estimate (95% CI) | Estimate (95% CI) | Estimate (95% CI) |
| 'full' 450k/27k | -0.28 (-1.03, 0.47) | 1.02 (0.96, 1.09) | 7.02 (6.25, 7.93) | 0.85 (0.84, 0.87) |
| 'reduced' 450k/27k | -0.32 (-1.09, 0.45) | 0.88 (0.81, 0.94) | 3.18 (2.40, 3.95) | 0.86 (0.85, 0.88) |

Regressions of DNAm age on chronological age within the full and reduced datasets, within each age group, reveal further age-dependent differences (Table 3). Among those <20 y, the slope in the reduced datasets is shallower and significantly differ (t-test, p=0.002) when compared with the full dataset, while the intercepts do not differ (t-test, p = 0.94). Among those ≥ 20 years, the slopes do not differ significantly (t-test, p= 0.84), but the underestimation of DNA methylation age by the reduced data, as compared to the full data, is 3.84 y (t-test, p<0.001) at the intercept.

### Potential impact of underestimation on regression outcomes

If the underestimation of DNA methylation age within each dataset is systematic, associations between DNAm age and clinical variable (or other variable of interest) in the reduced and full 450k datasets should be similar. Given the differences in DNAm age estimation for individuals age <20 y vs ≥20 y (Table 3, Figure S1), we examined associations between age acceleration and sex, (Table 4) in both age groups. Using DNAm age acceleration, the residuals of age regressed on DNAm age, the effect estimates obtained in the full 450k data were not significantly different from those obtained in the reduced 450k data in subjects aged 20 years or more (p = 0.87) nor in subjects <20 years (p = 0.22). This finding did not differ when we used epigenetic age in place of the age acceleration measure (not shown), and did not differ depending on whether the data was derived from the 27k array or 450k array. Residual violin plots for regressions of epigenetic age on sex (Figure S2) show no large or systematic differences in the distribution of epigenetic age residuals, further reinforcing the similarity of the regressions with and without the removal of the 17 probes missing from the 850k platform.

**Table 4.** Regressions of age acceleration on sex for CATHGEN450k and GEO datasets, using DNA methylation age calculated using the (full) 450k data and reduced 450k data. Regressions were conducted for each dataset individually, and then in aggregate while stratifying for chronological age (<20y and ≥20y). P-values result from a t-test to compare the slopes for regressions using the various DNAm ages.

| Dataset | N (prop. female) | <u>(Full) 450k/27k data</u> | <u>Reduced 450k/27k data</u> | <u>Full vs. reduced 450k/27k data</u> |
| --- | --- | --- | --- | --- |
|  |  | <u>DNAm age</u><br>Slope Est. (95%CI) | <u>DNAm age</u><br>Slope Est. (95%CI) | p value |
| Cathgen450k | 205 (0.38) | 0.28 (-1.31, 1.87) | 0.39 (-1.24, 2.02) | 0.92 |
| GSE20067 | 192 (0.51) | 0.03 (-1.64, 1.7) | 0.12 (-1.55, 1.8) | 0.94 |
| GSE20242 | 50 (0.74) | -3.31 (-17.88, 11.27) | -1.58 (-16.68, 13.51) | 0.87 |
| GSE32149 | 48 (0.52) | 1.89 (-1.58, 5.37) | 2.25 (-1.45, 5.96) | 0.89 |
| GSE40279 | 656 (0.52) | 1.41 (0.46, 2.36) | 1.39 (0.45, 2.33) | 0.98 |
| GSE41037 | 720 (0.38) | 1.25 (0.53, 1.97) | 1.04 (0.36, 1.73) | 0.68 |
| GSE41169 | 95 (0.29) | -0.89 (-2.57, 0.78) | -1.13 (-2.72, 0.46) | 0.84 |
| GSE42861 | 689 (0.71) | 0.17 (-0.69, 1.03) | 0.01 (-0.85, 0.87) | 0.80 |
| less than 20y | 662 (0.06) | -0.6 (-1.52, 0.32) | 0.19 (-0.69, 1.08) | 0.22 |
| 20y or older | 3,294 (0.60) | 1.51 (1.01, 2.01) | 1.57 (1.07, 2.07) | 0.87 |

## Discussion

Estimation of DNAm age is a methylation array dependent procedure, in so much as differing arrays may not have all probes used to develop the DNAm age estimator. Use of the epigenetic clock to estimate DNAm age from data generated from the Illumina MethylationEPIC array is likely to produce substantial underestimation of DNAm age, relative to the DNAm age estimated with the Illumina 450K array. A 3.3-year and 5-year increased DNAm age using the Horvath epigenetic clock has been associated with an increase of 10 body mass index units (19) and a 16% increase in mortality (20), respectively. Thus, observed underestimations, in the range of 4 years, could cause substantial mis-estimations of mortality and obesity risk based on the measured DNAm age if array differences are not accounted for. Using age-adjusted residuals (DNA methylation age acceleration) or adjusting for age when using Δage (DNAm age – chronological age) as a predictor since the correlation between chronological age and DNA methylation age appears to be independent of array. Systematic differences due to array design would alter the intercept in such models but not regression coefficients. Thus, regression models will reflect highly concordant results across arrays, but this will not necessarily be reflected in comparisons of absolute epigenetic aging differences with outcomes across methylation platforms. Estimating epigenetic age on a “reduced” 450k dataset (i.e. using probes only available on the 850k array) produced similar underestimation as observed when using the 850k data, indicating that the observed underestimation is primarily driven by the missing probes (Table 1), as opposed to technological differences between the 850k and the 450k arrays. This might be expected given the fact that the probes used for the 850k array used the same chemistry and color channels as previous probes.

This study employed many of the same publicly available GEO datasets used to develop the 450k clock, allowing direct comparisons in datasets which have been previously shown to estimate DNAm age well (1). We focused on blood, since that is the tissue for which the Horvath epigenetic age estimator provides the most accurate and consistent associations, and in which the Horvath DNAm age estimator has been most widely applied. Because CATHGEN 450k and 850k data were estimated on independent (i.e., non-overlapping) groups of individuals, direct comparison of the underestimation of DNAm age within individuals was not possible. However, the size of the CATHGEN datasets still offer the ability to compare these measures in the same source population, and both datasets were similar in age and sex makeup (Table 1).

The Illumina MethylationEPIC array represents a substantial step forward in the genome-wide assessment of DNA methylation. As DNA methylation array technology has progressed, researchers may wish to combine epigenetic age derived from 450k/27/k and 850k data; however, the deviation in DNAm age estimates among the array platform generations may introduce error into subsequent analyses. Thus, care should be taken when using epigenetic biomarkers, such as Horvath’s clock, that were developed using 450k and 27k data, as they may not be fully optimized for the Illumina MethylationEPIC array.

## Author contributions

RD and CWC are responsible for conception and design. RD was responsible for collecting, processing and analyzing the publicly available data. RD and LK carried out processing and analyses for CATHGEN. KO, DDS, RBD and WC provided funding for the creation of the epigenetic data for CATHGEN on the 850K platform, and provided guidance in design and execution of analyses. CH, ERH, SG, SS and WK are responsible for recruiting and maintaining the CATHGEN biorepository; they supplied both the demographic and epigenetic data for CATHGEN on the 450k platform. All authors have been involved in the editing of this paper and have reviewed the final draft.

## Acknowledgements

The authors would like to thank Kristen Rappazzo for critiques of the manuscript and analysis.

## Supporting Information

***Table S1. Summary of GEO datasets.***

***Figure S1. Plot of reduced 450k DNA methylation age by 450k data DNA methylation age in CATHGEN 450k data and the publicly available datasets for (a) all observations, (b) those < 20 years of age, and (c) those ≥ 20 years of age.** As can be seen across the plots, although the slope between the full and reduced DNA methylation age differs between the two age groups the overall correlation remains high.*

***Figure S2. Violin plots of residuals by sex, from regression of DNA methylation age acceleration on sex for 450k data, reduced 450k data, in the CATHGEN 450k and publicly available GEO datasets.** The distribution of residuals from the regression of age acceleration on sex is the same even after removing the 17 probes, indicating that regressions using age acceleration from the reduced 450k data (which underestimates DNA methylation age) remain valid as the underestimation is captured as an intercept shift in the models.*

